# Analysis of human brain tissue derived from DBS surgery

**DOI:** 10.1101/2021.06.18.448926

**Authors:** Salla M. Kangas, Jaakko Teppo, Maija J. Lahtinen, Anu Suoranta, Bishwa Ghimire, Pirkko Mattila, Johanna Uusimaa, Markku Varjosalo, Jani Katisko, Reetta Hinttala

## Abstract

**Background:** Transcriptomic and proteomic profiling of human brain tissue is hindered by availability of fresh samples from living patients. Postmortem samples usually represent the advanced disease stage of the patient. Furthermore, the postmortem interval affects the observed transcriptomic and proteomic profiles. Therefore, access to fresh brain tissue samples from living patients is valuable resource to obtain information on metabolically intact tissue. The implantation of deep brain stimulation (DBS) electrodes into the human brain is a neurosurgical treatment for, e.g., movement disorders. Here, we describe an improved approach to collect brain tissue from surgical instruments used in implantation of DBS device for transcriptomics and proteomics analyses.

**Methods:** Samples were extracted from guide tubes and recording electrodes used in routine DBS implantation procedure that was carried out to treat patients with Parkinson’s disease, genetic dystonia and tremor. RNA sequencing was carried out to tissue extracted from the recording microelectrodes and liquid chromatography-mass spectrometry was carried out to analyze tissue from guide tubes. To assess the performance of the current approach, obtained datasets were compared with previously published datasets representing brain tissue.

**Results:** In RNA sequencing, altogether 32,034 transcripts representing unique Ensembl gene identifiers were detected from eight samples representing both hemispheres of four patients. By using liquid chromatography-mass spectrometry, we identified 734 unique proteins from 31 samples collected from 14 patients. Comparison with previously published brain derived data indicated that both of our datasets reflected the expected brain tissue specific features. The datasets are available via BioStudies database (accession number S-BSST667).

**Conclusions:** Surgical instruments used in DBS installation retain enough brain material for protein and gene expression studies. Analysis of the datasets indicated that hemisphere-specific expression data can be obtained from individual patients without any sample pooling and without any modifications to the standard surgical protocol. Comparison with previously published datasets obtained with similar approach proved the robustness and reproducibility of the current improved protocol. This approach overcomes the issues that arise from using postmortem tissue, such as effect of postmortem interval, on proteomic and transcriptomic landscape of the brain and can be used for studying molecular aspects of DBS-treatable diseases.

## Background

Neurodegenerative diseases, especially Alzheimer’s and Parkinson’s, are widely studied using postmortem brain in the search for disease biomarkers and for understanding of the molecular basis of the disease (1–5). When using postmortem samples, the integrity of brain tissue is compromised due to the delay in collecting the samples, which may bias the results. Some proteins are more prone to degradation than others, and the observed outcome may depend on postmortem interval (5,6). Likewise, RNA is rapidly degraded (7,8), and fresh human brain transcriptome differs from postmortem transcriptome essentially (9). Therefore, access to fresh brain tissue is critical in obtaining accurate information about brain-specific transcripts and the transcriptome *in vivo*. Recently, approaches that utilize fresh, non-tumorous brain-derived samples from patients treated for various brain-affecting conditions have emerged. For example, brain biopsy samples were collected from patients suffering traumatic brain injury in conjunction with the insertion of an intracranial pressure-monitoring device during corticotomy (10).

Deep brain stimulation (DBS) is a neurosurgical treatment for advanced and medically refractory movement disorders, such as Parkinson’s disease, essential tremor and dystonia. In addition, pain, epilepsy and psychiatric disorders are increasingly treated with DBS (11). During DBS operation, intracranial electrodes are targeted into specific locations in the deep brain structures bilaterally. The intracranial leads are connected to an external impulse generator through extension leads. The DBS device stimulates deep brain structures with a low-level electrical current that alleviates patients’ symptoms in a reversible manner. Zaccaria *et al*. have previously collected brain-derived material during DBS surgery from several individual patients with Parkinson’s disease for proteome and transcriptome analysis (12). Their approach included additional step during the surgery, where a blunt stylet was inserted into the brain tissue to collect the material for analysis. Here, our aim was to assess whether the surgical, non-permanent instruments used in the standard DBS implantation procedure as followed at the Operative Care Unit at Oulu University Hospital (13) contain enough hemisphere-specific brain-derived material from individual patients for RNA sequencing (RNA-seq) and liquid chromatography-mass spectrometry (LC-MS).

## Results

Here, we demonstrate the collection of fresh brain tissue samples from single-use surgical instruments, guide tubes and recording microelectrodes, which are needed for the DBS electrode implantation into movement disorder patients, and the omics datasets derived from subsequent analyses of the tissue material at the RNA and protein levels (Figure 1A).

**Figure 1.**
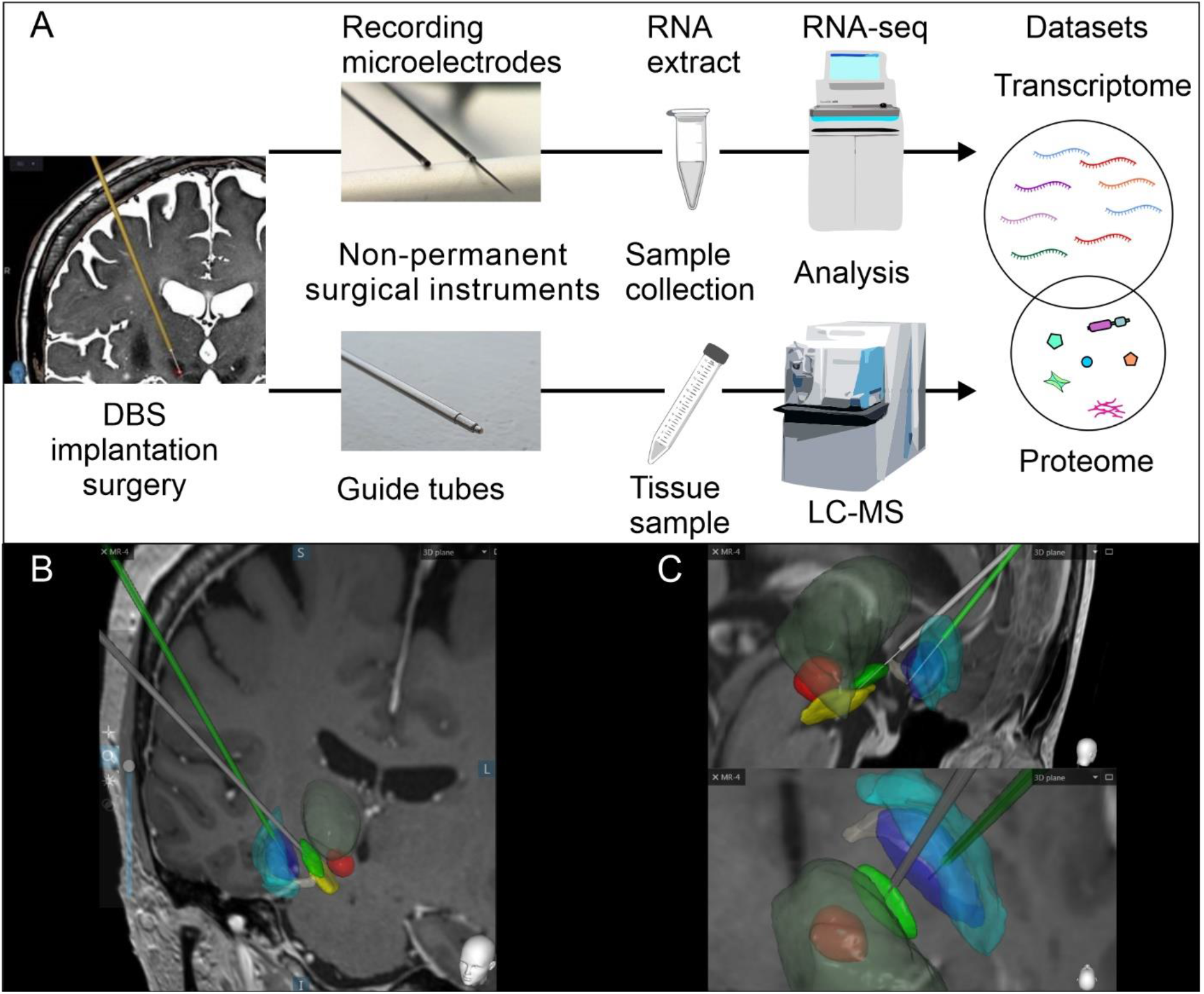
Workflow to collect fresh brain material during DBS surgery for treatment of patients with movement disorders. A) DBS leads were implanted into patients to treat movement disorders in neurosurgical operation at Operative Care Unit, Oulu University Hospital, and the samples were collected from the guide tubes from both hemispheres of 14 patients for LC-MS analysis and the recording microelectrodes of four patients for RNA-seq analysis during the standard DBS implantation procedure. The tissue samples for LC-MS were collected from the guide tubes that protruded through the brain tissue to reach the target area and therefore contained tissue material from different brain regions. The RNA samples for sequencing were collected from the recording microelectrodes targeted to the subthalamic nucleus (STN) and globus pallidus interna (GPi). B) The image shows how the guide tubes (grey and green thick lines) passed through brain tissue and the most distal end was 10 mm from the planned target. C) In contrast, the microelectrodes (thin grey lines) travel inside the guide tube, and they touched brain tissue only in the STN (green area) and GPi (blue area). To help with anatomical orientation, the other brain structures are the thalamus (dark transparent green), substantia nigra (yellow), red nucleus (red), ansa lenticularis (dark white) and globus pallidus externa (transparent turqoise).

The samples were collected from 17 patients treated with DBS at the Operative Care Unit at Oulu University Hospital for RNA-seq (n=4) and LC-MS (n=14) (Tables 1 and 2, respectively).

**Table 1.**
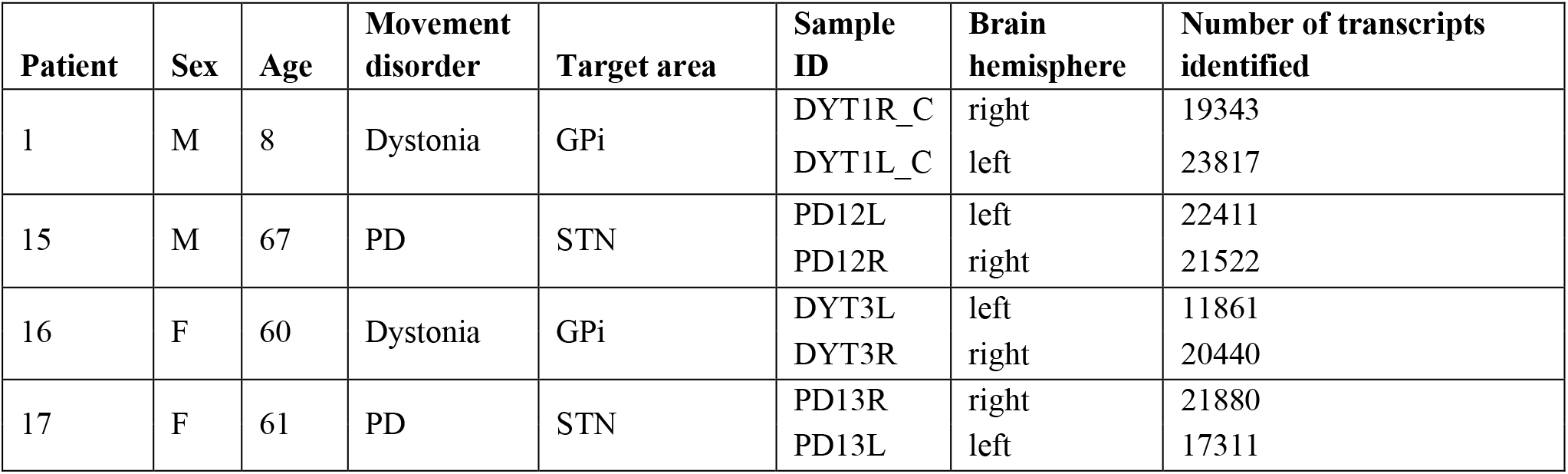
Info about patients and samples for RNA-seq analysis.

**Table 2.** Info about patients and samples for LC-MS analysis.

| Patient | Sex | Age | Movement disorder | DBS target area | Sample ID | Brain hemisphere | Visible blood | Total protein (µg) | Number of proteins identified |
| --- | --- | --- | --- | --- | --- | --- | --- | --- | --- |
| 1 | M | 6 | Dystonia | GPi | DYT1L_A | left | yes | 453.11 | 181 |
|  |  |  |  |  | DYT1R_A | right | no | 4.28 | 298 |
| 2 | M | 54 | Tremor | VIM | TRE1R | right | no | 8.51 | 217 |
|  |  |  |  |  | TRE1L | left | no | NA | 190 |
| 3 | M | 67 | PD | STN | PD1L | left | no | NA | 165 |
|  |  |  |  |  | PD1R1 | right | no | 14.25 | 375 |
|  |  |  |  |  | PD1R2 | right | no | NA | 218 |
| 4 | F | 58 | PD | STN | PD2R | right | yes | 82.63 | 199 |
|  |  |  |  |  | PD2L | left | no | 6.03 | 319 |
| 5 | M | 58 | PD | STN | PD3L | left | yes | 86.80 | 153 |
|  |  |  |  |  | PD3R | right | yes | 148.61 | 252 |
| 6 | M | 62 | PD | STN | PD4L | left | yes | 63.82 | 211 |
|  |  |  |  |  | PD4R | right | yes | 93.01 | 175 |
| 7 | M | 59 | PD | STN | PD5R | right | no | 2.36 | 244 |
|  |  |  |  |  | PD5L | left | yes | 87.60 | 262 |
| 8 | F | 67 | PD | STN | PD6R | right | no | NA | 292 |
|  |  |  |  |  | PD6L | left | yes | 20.07 | 161 |
| 9 | M | 59 | PD | STN | PD7L | left | no | 3.87 | 136 |
|  |  |  |  |  | PD7R | right | no | NA | 150 |
| 10 | M | 52 | PD | STN | PD8R | right | no | NA | 193 |
|  |  |  |  |  | PD8L | left | no | NA | 67 |
| 11 | F | 63 | Dystonia | GPi | DYT2L | left | no | NA | 120 |
|  |  |  |  |  | DYT2R | right | no | NA | 200 |
| 12 | M | 59 | PD | STN | PD9L | left | no | NA | 178 |
|  |  |  |  |  | PD9R | right | yes | 9.24 | 261 |
| 13 | F | 66 | PD | STN | PD10L | left | yes | 14.79 | 223 |
|  |  |  |  |  | PD10R | right | no | 2.51 | 211 |
| 1 | M | 7 | Dystonia | GPi | DYT1L_B | left | yes | 50.52 | 163 |
|  |  |  |  |  | DYT1R_B | right | yes | 48.88 | 287 |
| 14 | F | 62 | PD | STN | PD11R | right | no | NA | 169 |
|  |  |  |  |  | PD11L | left | no | NA | 204 |

### RNA sequencing analysis produces tissue-specific data

Transcriptomic analysis was focused on the subthalamic nucleus (STN) or globus pallidus interna (GPi) regions, which are specific targets of DBS in treating patients with movement disorders (Figure 1C). Samples for RNA-seq were collected separately from the recording microelectrodes targeted to both hemispheres of four patients (Table 1), of whom two had Parkinson’s disease and STN as the target area and two had genetic dystonia and GPi as the target area. The number of identified genes expressed in the eight samples varied from 11,861 to 23,817 (Figure 2A), of which 32,034 genes were unique across all the samples (Additional file 1). In total, 14,562 genes identified were shared between all samples from the STN (Figure 2B), and 10,638 genes identified were shared between all samples from the GPi (Figure 2C). Also, 9,901 genes were commonly detected in all samples from both STN and GPi regions.

**Figure 2.**
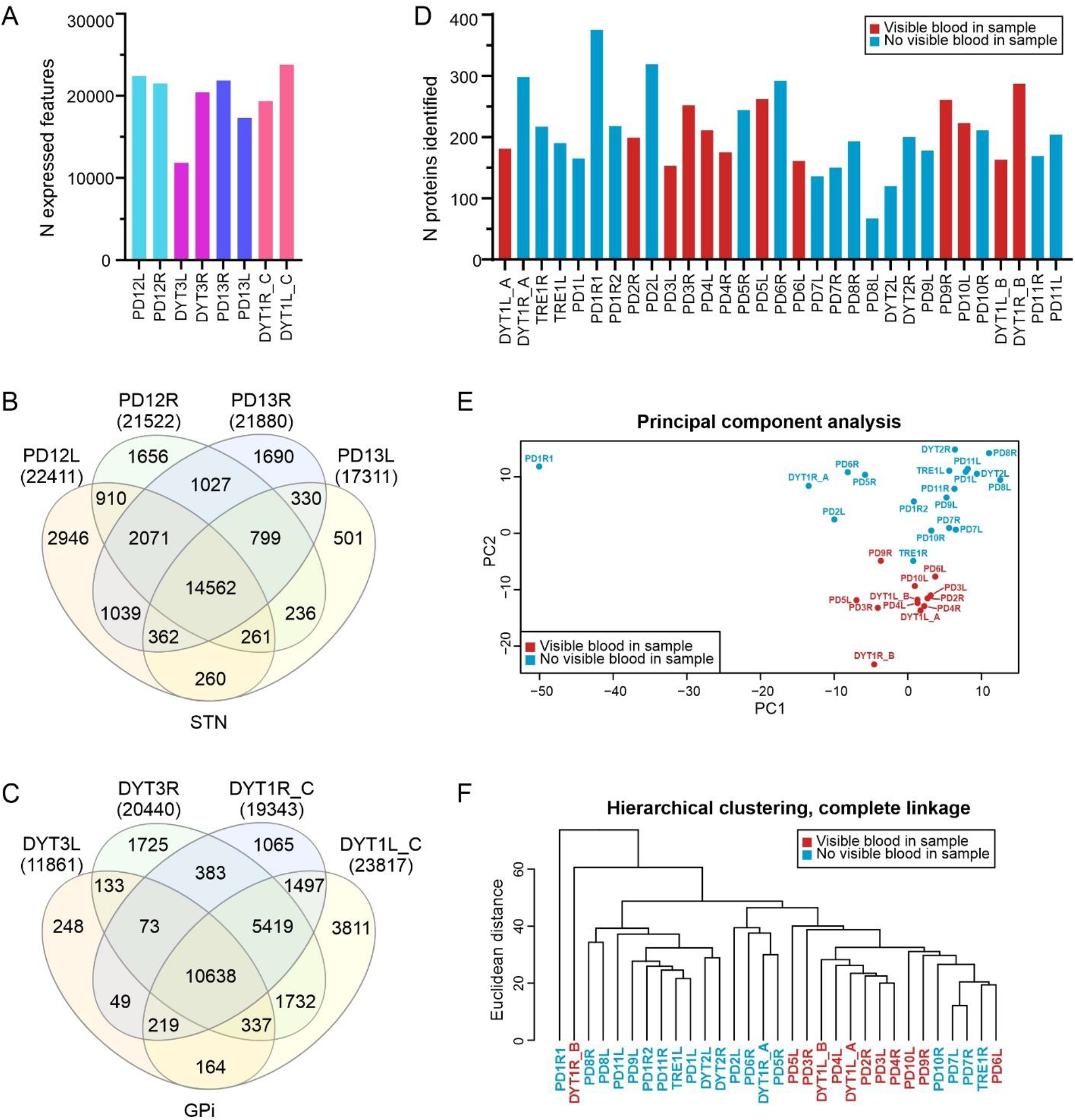
Features of proteomics and transcriptomics datasets obtained from the RNA sequencing and LC-MS analyses of the patient-derived brain tissue. The sample encoding indicates the patients’ disorders are as follows: Parkinson’s disease (PD, n=13), genetic dystonia (DYT, n=3) and tremor (TRE, n=1). A) The number of expressed genes in each sample. Venn diagrams show the number of common genes identified in the samples from B) the subthalamic nucleus (STN) and C) the globus pallidus interna (GPi) target areas. D) The number of identified proteins in each sample, colored based on whether blood was visible in the sample. No statistical difference in the number of proteins identified was observed (t-test p=0.51). E) Principal component analysis (PCA) plot of the proteomic data and F) hierarchical clustering, colored based on whether blood was visible in the sample, shows that samples with visible blood tend to cluster.

### LC-MS can be used to analyze the brain hemisphere-specific tissue samples attached to the guide tubes

Proteomics analysis was carried out from the tissue material that had attached to the guide tubes that are used to reach the microelectrode target region (Figure 1B). After confirming by immunoblotting that brain-specific proteins were present in the tissue material collected from guide tubes (Additional file 2), we proceeded with proteomics analysis. By using LC-MS, we could identify 734 unique proteins from 31 samples from 14 patients (Table 2). Eighteen of these proteins (seven being abundant in blood) were present and quantified in all samples (Additional file 3). Based on visual inspection, the samples contained variable amounts of blood, which did not influence on the overall number of proteins identified in those specific samples (Figure 2D). However, the clustering of the samples according to the blood observed in them was evident (Figures 2E and 2F). When we analyzed the identified protein datasets using DAVID (14,15), we found that the enriched GO terms reflected the brain tissue well, and blood, which was observed in some of the samples, was not over-represented among these terms (Figure 3F, Additional file 3). However, we believe that the removal of blood from the samples prior to the LC-MS analysis will improve the specificity of the assay, if managed with minimal sample loss. When the transcriptomics dataset was mapped to Uniprot identifiers, the overlap between the transcriptomics dataset and proteomics dataset was 686 identifiers, which covers 93,5% of identified proteins.

**Figure 3.**
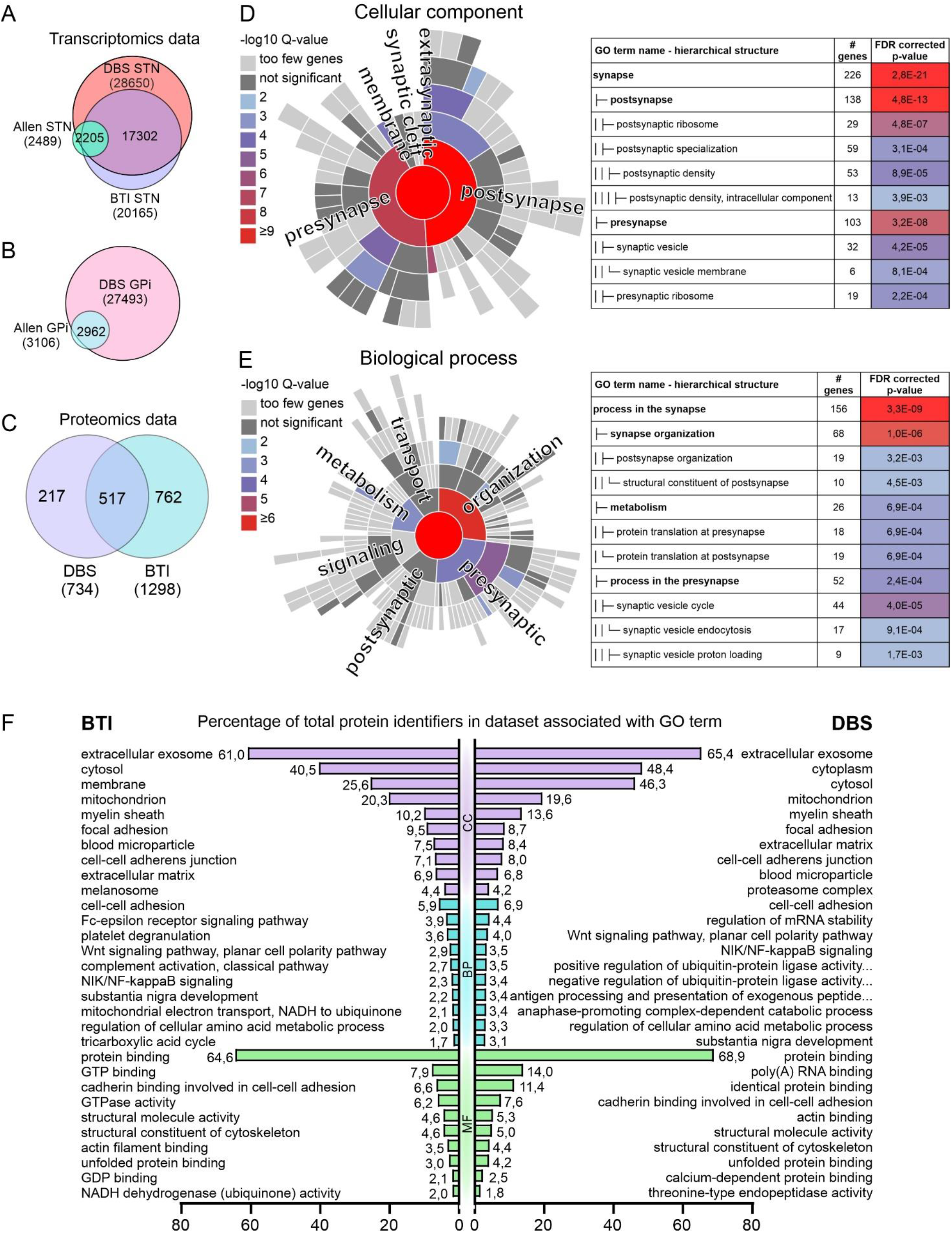
Comparison of the datasets to other published data and Gene Ontology (GO) enrichment analyses. We compared our target region-specific transcriptomics datasets to the anatomically specific expression datasets in Allen Brain Atlas and found substantial overlap in A) STN- and B) GPi-specific terms. The BTI dataset contained only STN data, and we found that our data had 86% overlap with the BTI dataset. C) We compared our dataset of all unique protein identifiers to the BTI proteomics dataset(12) and found that 517 identifiers out of 734 (70%) were in common with the BTI dataset. We tested the top 20% expressed RNA-seq identifiers common to all analysed samples (n=1,980) using the SynGO Knowledge base gene set enrichment tool(17). D) Ten terms in the cellular component category and E) eleven terms in the bioprocess category were significantly enriched at 1% FDR (testing terms with at least three matching input genes). F) To obtain an overview of the type of proteins identified, GO enrichment analysis using DAVID bioinformatics platform Version 6.8 was performed to the list of all identified proteins across all the samples. We also performed the same analysis to the BTI dataset published previously by Zaccaria *et al*.(12) to compare the outcomes of these two methods. The top 10 most enriched terms in each GO category (Cellular component, CC; Biological process, BP; Molecular function, MF) show that our dataset shares many top terms with the BTI dataset.

### Comparison between the current approach and the previously published method by Zaccaria *et al*

Zaccaria *et al*. have previously utilized DBS surgery to obtain brain-derived material, which they termed “brain tissue imprints” (BTIs), for proteome and transcriptome analysis from Parkinson’s disease patients (12). To our knowledge, this is the only published method that resembles ours; however, there are substantial differences in procedure. We compared our approach and results to the approach published earlier by Zaccaria *et al*. (Table 3) (12).

**Table 3.** Comparison of the two approaches to collect and analyze samples obtained during DBS surgical procedure.

|  | **Sample collection procedure described in the current paper** | **BTI method (12)** |
| --- | --- | --- |
| **Sample collection protocol** | The brain tissue attached to the guide tubes and microelectrodes during standard DBS implantation procedure were used for sample preparation. | A blunt stylet was inserted through the guide tube into the brain tissue during DBS implantation procedure for one minute to obtain material for analyses. |
| **Sample usage** | Both RNA-seq and LC-MS analyses can be carried out from the same individual patients and their separate brain hemispheres (if patient is awake during the procedure). | Tissue sample was used either for RNA microarray analysis or pooled for Nano-LC-MS/MS. Samples were also alternatively used for immunocytochemistry or scanning electron microscopy. |
| **Transcriptomics** | The tissue material was collected from the recording microelectrode targeted to the specific well-defined area in the deep brain region and it was prepared for RNA-seq. | RNA microarray from the tissue sample attached to the blunt stylet was carried out after application of double amplification protocol. |
| **Proteomics** | No pooling of samples. Hemisphere-specific LC-MS data was obtained from the tissue collected from the guide tube. | Six samples from different patients and brain hemispheres were pooled for in-gel fractionation and subsequent MS analysis. |

Zaccaria *et al*. collected 19 samples from twelve patients as follows: After determining the DBS target area via MER, a blunt stylet, as an additional sample collection step, was inserted through the guide tube into the brain for one minute to obtain material for analyses. The material attached to the stylet was then used for proteomics, electron microscopy, immunohistochemistry and immunofluorescence or RNA microarray analysis. In our protocol, no alterations or additional steps were introduced to the standard DBS procedure; instead, we collected the brain tissue material that had attached to the guide tubes and recording microelectrodes used during the normal surgical procedure. Because our DBS implantation surgery followed standard procedure, we were able to collect samples systematically from both hemispheres of each patient, whereas the protocol used by Zaccaria *et al*. had technical constraints that allowed sample collection procedure from both hemispheres only occasionally. Essential difference in the transcriptomics data output is that Zaccaria *et al*. did RNA microarray analysis whereas we did RNA sequencing. RNA microarray analysis profiles predefined transcripts through hybridization while RNA sequencing is both quantitative and it covers whole transcriptome and therefore it can be used to detect different not-predefined transcripts. During proteomics analysis, Zaccaria *et al*. pooled multiple samples, while we analyzed hemisphere-specific individual samples.

### Comparison of the datasets to the previously published data

The datasets achieved via our method were compared to the BTI approach previously described by Zaccaria *et al*. (12). In their transcriptomics analysis, Zaccaria *et al*. produced three RNA microarray datasets from the STNs of three patients, containing 35,701; 29,842 and 27,350 detected unique microarray probe identifiers. We converted the probe identifiers from the BTI RNA microarray dataset into 20,165 unique Ensembl gene (ENSG) identifiers to allow comparison with our dataset. Ultimately, 17,302 unique identifiers (86%) were common between our STN-specific RNA-seq and BTI microarray datasets (Figure 3A). We also compared our STN- and GPi-specific RNA-seq datasets to brain region-specific expression datasets representing up-regulated genes in the STN (Figure 3A) and GPi (Figure 3B) (Allen brain atlas) and found that a substantial majority (85% and 95%, respectively) of the upregulated genes were present in our datasets. The BTI protocol for sample acquisition led to the identification of 1,298 unique proteins from the samples using Nano-LC-MS/MS (12). We compared the list of our identified proteins to the BTI proteomics dataset and found that 70% of the proteins in our dataset are common with the BTI proteomics dataset (Figure 3C).

SynGO is a knowledgebase that focuses on synapse-specific ontologies, and its annotations are based on published, expert-curated evidence (17). In their study, Koopmans and others show that synaptic genes are exceptionally well conserved and less tolerant of mutations than other genes (17). They conclude that many SynGO terms are overrepresented for genes that have variants associated with brain disease. By using the SynGO analysis tool, we could identify several terms enriched among the 20% top of expressed genes (Figures 3D and 3E, Additional file 4), and 68% (754/1,112) of SynGO annotated genes were found in our RNA-seq dataset, which contained 9,901 genes overlapping all the eight samples. This indicates that the RNA-seq data obtained from the tissue attached to the recording microelectrodes during the DBS implantation procedure are a potentially useful resource in studying brain disorders and the brain-specific transcriptome landscape, such as brain-specific transcript isoforms *in vivo*.

The GO enrichment analysis of our current and the BTI proteomics datasets via DAVID revealed that both datasets have many similar GO term profiles in their top 10 enriched terms (Figure 3F). For their analysis, Zaccaria *et al*. pooled samples from six patients and brain hemispheres for in-gel fractionation and subsequent MS analysis, whereas our data are both patient- and hemisphere-specific (12). In general, sample pooling increases the number of proteins identified, but it also leads to the loss of information on sample variation, the missed detection of biomarkers and the false identification of others (18). Molinari *et al*. found that pooled samples are not equivalent to average of biological values and pooling affects statistical analysis (18). The pooling of the BTI samples (12) for downstream analyses has led to the loss of substantial patient- and hemisphere-specific information, whereas our datasets are patient- and hemisphere-specific.

## Discussion

Published proteomics and transcriptomics datasets for different human brain areas are most often based on postmortem material because brain biopsies from living patients are hardly achievable. When using postmortem samples, the integrity of brain tissue is compromised due to the delay in collecting the samples, which may bias the results. Dachet *et al*. showed that, during the postmortem interval, within few hours, neuronal gene expression, especially in the case of brain activity-dependent genes, declines rapidly, while astroglial and microglial gene expression increase reciprocally (9). In turn, most of the housekeeping genes, which are frequently used for normalization in the comparison of expression levels, are very stable. Also, reduced diversity in the complexity of differentially spliced transcripts in the case of the ultra-complex splicing pattern of RBFOX1 was demonstrated (9). Using fresh material that is processed rapidly within a known time window reduces the technical variation caused by postmortem changes. Biopsies from brain tumors, such as gliomas, are one source of fresh brain-derived tissue that has been utilized quite widely in various omics approaches during past years (19,20), even though they represent the neoplastic phenotype, which does not correspond to normal brain tissue.

Our results indicate that the approach described here to collect patient-derived fresh brain tissue from instruments used for DBS implantation surgery is applicable in studying brain disorders at the individual hemisphere level. The collection of defined patient cohorts and comparisons of the disease-specific brain proteomes and transcriptomes are a valuable tool for use in identifying disease signatures. Compared to postmortem samples, DBS implantation-derived samples present earlier time points in the disease course and phenotype, which helps in understanding the changes that occur at defined clinical stages during the development of neurological symptoms. One potential caveat regarding this approach is a lack of healthy controls, but a comparison of the molecular signatures between different diseases and stages of disease progression allows the identification of common brain-specific proteoforms and transcripts, as well as novel disease-specific markers, for further studies. If a neurosurgical operation is performed on a conscious patient to perform MER and thus adjust the target region, at the same time, samples for transcriptomics can also be collected from the microelectrodes representing a very defined brain target area. In contrast to this highly region-specific transcriptomics analysis, our proteomics analysis provides data from a cross-section of the brain, containing an expression profile from a mixture of cell types from different brain layers.

As essential improvements to the BTI approach previously described by Zaccaria *et al*. (12), our method does not make any modifications to our standard surgical DBS procedures, and our approach allows collecting samples from the guide tubes from both hemispheres of the patients routinely, without sample pooling for subsequent analyses. Pooling of samples may increase the number of identified proteins, but at the same time, it masks sample-specific proteoforms and post-translational modifications and valuable information on individual patients is lost (18). Different posttranslational modifications may reflect disease stage and thus function as disease (stage) biomarkers. Comparison between the datasets generated by the BTI approach by Zaccaria *et al* and by us confirmed that the approach in general is reproducible and robust despite that we have differences in sample collection procedure, and we have used different analysis platforms. Both transcriptomics and proteomics datasets contained substantial number of common identifiers.

## Conclusions

The improved approach we describe here can be used to bring novel information about brain tissue-specific transcript variants from specific brain regions, proteoforms and post-translational modifications, which represents valuable additional knowledge about the brain transcriptome and proteome landscape *in vivo*. Analysis of fresh brain material is important because postmortem changes lead to a rapid loss of transcriptomic diversity in neuronal tissue and glial activation causes the upregulation of gene expression that does not correspond the normal expression landscape of the brain tissue (9). In the future, as proteomics and transcriptomics techniques become more sensitive and new methods are developed, the approach described here will be a valuable tool with which to access fresh brain-derived material for novel discoveries. By combining the patient-derived proteomics and transcriptomics data with experiments utilizing patient-derived cells and disease modelling, this approach advances personalized medicine and studies in the field of neurological diseases.

## Methods

### Patients

DBS leads were implanted into patients during neurosurgical operations at the operative care unit, Oulu University Hospital, Finland, between October 2017 and June 2019. The indications for DBS treatment were Parkinson’s disease (n=13), genetic dystonia (n=3) and tremor (n=1). Guide tubes and microelectrodes were used during the standard DBS implantation procedure (Figure 1). The samples were collected from the recording microelectrodes collected from four patients (eight samples) for RNA sequencing (RNA-seq) (Table 1) and guide tubes from 14 patients (31 samples) for liquid chromatography-mass spectrometry (LC-MS) (Table 2). When extracting the RNA from the recording microelectrodes, the target region was the STN (n(patients)=2, n(samples)=4) and GPi (n(patients)=2, n(samples)=4). When collecting the tissue from the guide tubes, intracranial leads were targeted into the subthalamic nucleus (STN, n(patients)=11, n(samples)=23), globus pallidus interna (GPi, n(patients)=2, n(samples)=6), or ventral intermediate nucleus of the thalamus (VIM, n(patients)=1, n(samples)=2). The LC-MS samples from Patients 4 and 6 (sample codes PD2 and PD4, respectively) were obtained during re-implantation to resume DBS treatment after the previous removal of the DBS leads due to technical failure. Two samples, one from each hemisphere (coded L=left or R=right), were obtained from each patient during the procedure, with one exception. Three guide tubes, of which two (PD1R1 and PD1R2) were from the right hemisphere, were obtained from Patient 3.

From Patient 1, three sets of samples were obtained from three separate surgical procedures. The first samples (DYT1L_A and DYT1R_A) were collected from the guide tubes for LC-MS analysis during the first DBS implantation procedure. The second samples were collected for LC-MS analysis during revision surgery, which was performed due to technical failure. The brain tissue samples were collected from the revised DBS leads. The third samples (DYT3L_A and DYT3R_A) were collected for RNA-seq from the recording microelectrodes during the reimplantation of the DBS.

### DBS implantation procedure

The surgical procedure for DBS implantation was carried out according to the standard protocol in our institute, as described in detail by Lahtinen *et al*. (13). The patient-specific targeting of intracranial electrodes was planned on brain magnetic resonance images (MRI) and adjusted with intraoperative clinical testing and microelectrode registration (MER) during neurosurgical operation if the patient was awake. The guide tubes (Universal Guide Tube, Elekta, Stockholm, Sweden) and recording microelectrodes (Leadpoint, Alpine Biomed, Skovlunde, Denmark) used during the implantation procedure were collected and used for sample acquisition for proteomics and transcriptomics, respectively. The location of the intracranial electrodes is most commonly in the deep basal nuclei, and the most common trajectory to the target area is through the posterior parts of the frontal lobes (Figure 1B-C).

### RNA extraction for RNA-seq

After the removal from the brain, the recording microelectrode was taken to a research laboratory on ice, where it was immediately immersed in 700 µl of QIAzol Lysis Reagent (Qiagen) at room temperature and triturated by using lead as a piston. The microelectrode was kept in QIAzol Lysis Reagent for about 10 minutes and triturated once more before discarding the electrode. The sample was briefly vortexed (2–3s) and stored at -80C. RNA-seq was performed by the sequencing unit of the Institute for Molecular Medicine Finland FIMM Technology Centre, University of Helsinki.

### RNA-seq

Total RNA was extracted with a Qiagen miRNeasy micro kit (QIAGEN, Hilden, Germany), according to the kit handbook. The quality and quantity of the extracted RNA samples were analyzed with a 2100 Bioanalyzer using an RNA 6000 Pico Kit (Agilent, Santa Clara, CA, USA). Paired-end cDNA libraries were prepared from 0,2 ng of extracted RNA, with eleven cycles of amplification using a SMART-Seq v4 Ultra Low Input RNA Kit, according to the manufacturer’s user manual (Takara Bio USA, Inc. Mountain View, CA, USA). One hundred pg of amplified cDNA was tagmented and indexed for sequencing using a Nextera XT DNA Library Prep Kit (Illumina, San Diego, CA, USA). LabChip GX Touch HT High Sensitivity assay (PerkinElmer, USA) was used for quality measurement and quantification of the purified dual-indexed libraries for equimolar pooling. The sequencing of the pooled samples was performed with an Illumina NovaSeq 6000 System (Illumina, San Diego, CA, USA). The read length for the paired-end run was 2×101 bp, and the target coverage was 15 M reads for each library.

### RNA-seq data analysis

The RNA-seq datasets were analyzed using FIMM-RNAseq data analysis pipeline Version v2.0.1. (Figure 4). The pipeline is implemented in Nextflow (21). Nextflow allows the portability and scalability of the pipeline and supports major cloud computing and batch processing technologies. More importantly, the pipeline allows the reproducibility of the results by using a version labeled set of software dependencies, a Conda environment. A Conda environment can be created manually, or Nextflow can be instructed to create one during a run-time without an effort. Alternatively, a readily available Docker image containing all software dependencies can be used to run the pipeline in a containerized computing environment, such as Docker and Singularity. Source code and a comprehensive user’s manual of the pipeline is available at https://version.helsinki.fi/fimm/fimm-rnaseq

**Figure 4.**
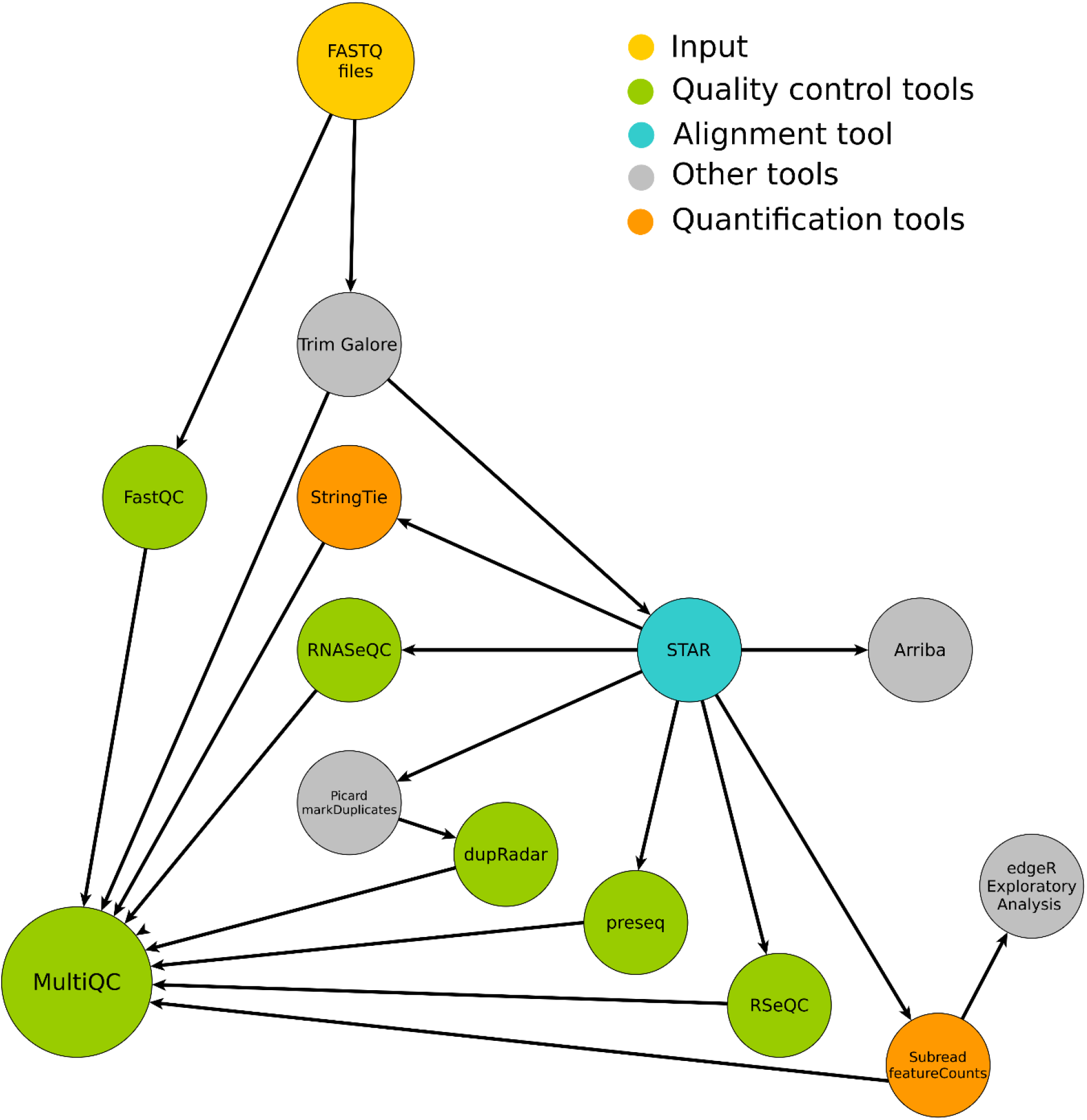
FIMM-RNAseq data analysis pipeline. FIMM-RNAseq incorporates quality control tools, such as FastQC and the pre-processing tool Trimgalore. It aligns RNA-seq reads using a STAR(32) aligner and performs gene quantification and transcript assembly using Subread(33) and StringTie(34), respectively. Extensive RNA-seq quality matrices are generated using RNASeQC(35), RseQC(36), dupRadar(37) and Preseq(38,39). An aggregated report from the major analysis steps is generated using MultiQC(40). Exploratory data analysis is performed using R and edgeR(41). As an optional component, the pipeline has the gene-fusion prediction tool Arriba(42).

### Brain tissue sample collection for proteomic analysis

Guide tubes were transported from the operation room to the laboratory on ice immediately after removal from the brain, and the samples were prepared for cryopreservation within one hour, as described below. The instruments from different hemispheres of each patient were handled individually.

Guide tubes were rinsed from inside with 10 ml of ice-cold phosphate-buffered saline (Sigma-Aldrich) using a 27-gauge needle and syringe. The suspension was collected into a 15-ml conical tube on ice. The tissue was pelleted via centrifugation at +4°C with 400 x g for 15 minutes. After centrifugation, the supernatant was removed carefully, and the pellet was flash frozen in liquid nitrogen. The samples were stored at -70°C until analysis.

### Sample preparation and LC-MS analysis

The cells were lysed, and the proteins denatured by adding 200 μL of 8 mol/L urea (Sigma-Aldrich), followed by 15 min sonication. Insoluble cell debris was removed via two rounds of centrifugation (15 min, 20817 G, 22 °C). Total protein content was measured with BCA assay (Thermo Scientific), the results of which are shown in Table 2.

Disulfide bonds were reduced with dithiothreitol (final concentration 5 mmol/L; Sigma-Aldrich), and the cysteine residues were carbamidomethylated with iodoacetamide (final concentration 15 mmol/L; Sigma-Aldrich), after which a pooled quality control (QC) sample was created by taking 31 μL of each sample and combining them. The proteins were digested with 2.5 μg of sequencing grade modified trypsin (Promega). The resulting peptides purified with C18 MicroSpin columns (The Nest Group, Inc.); for samples with > 60 μg of total protein, only 60 μg of total digested protein was taken for C18 purification, whereas for samples with < 60 μg of total protein, all of the sample was used. After C18 purification, the samples were evaporated to dryness with a vacuum centrifuge and stored at -20 °C.

Prior to LC-MS analysis, the samples were resolubilized with 15 min sonication in 30 μL of 1% acetonitrile + 0.1% trifluoroacetic acid in LC-MS grade water (all from VWR). The injection volume (between 2 and 10 μL) was determined based on the amount of total protein in the sample. The sample was injected into the LC-MS, separated with EASY-nLC 1000 (Thermo Scientific) using a 120 min linear gradient and detected with Orbitrap Elite MS (Thermo Scientific) using top20 data-dependent acquisition, in which the 20 most intense ions from each MS1 full scan are fragmented and analyzed in MS2. Pooled QC samples were analyzed at the beginning and end of the run sequence, but they were removed from the final data analysis.

Protein identification and quantification were performed with Andromeda and MaxQuant (22,23) with the standard settings and using a reviewed *Homo sapiens* UniProtKB/Swiss-Prot proteome (20431 entries, downloaded on 2019-08-30; The Uniprot Consortium (24)). In addition, LFQ (label-free quantification) was enabled, and identification FDR < 0.01 filtering was applied on both the peptide and protein levels. The LFQ intensity was used as an estimate of protein abundance without further normalization. From the output, we filtered decoy hits, proteins flagged as potential contaminants (but not serum albumin) and proteins identified with a modification site only. The LFQ intensities of all quantified proteins in all samples are presented in the Additional file 3. To account for the variable amounts of blood in the samples, the correlations of each protein’s LFQ intensity with those of serum albumin and hemoglobin subunit alpha were calculated, but no filtering based on these correlations was applied. The correlations are listed in the Additional file 3.

### Bioinformatics

To compare the overlap between the published datasets, the g:Convert tool of the g:Profiler (25) web server was used to convert the identifiers to the same namespace (ENSG_ID).

Allen Brain Atlas Adult Human Brain Tissue Gene Expression Profiles (16) for the subthalamic nucleus and globus pallidus internal segment were downloaded from https://maayanlab.cloud/Harmonizome/dataset/Allen+Brain+Atlas+Adult+Human+Brain+Tissue+Gene+Expression+Profiles. The reference list was formed by including all the upregulated genes from both hemispheres of the anatomical structure to the same list.

The SynGO portal (17) was used to analyze the enriched terms in the RNA-seq dataset. The used dataset was prepared for analysis by using the gene list that contained the overlapping genes among all RNA-seq samples (n=9901). To make a list of the top 20% of expressed genes among this common gene set, the expression levels of individual genes were normalized against the total expression level of the sample. The average value of normalized expression levels was used to rank the genes according to their expression level from high to low, and top 20% (n=1980) identifiers were used for SynGO analysis. A list of the ranked genes and original SynGO results appear in Additional file 4.

Gene ontology (GO) (26,27) enrichment analysis using DAVID (14,15) bioinformatics platform Version 6.8 was performed for the list of all identified proteins across all samples. The complete list of all enriched terms (FDR < 0.01) appears in Additional file 3. Both collapsed (DIR) GO terms and the uncollapsed (ALL) GO terms are listed in Additional file 3.

BioVenn (28) was used to draft area-proportional Venn diagrams. InteractiVenn was used to draw other Venn diagrams (29).

## Additional files

***Additional file 1:*** RNA-seq dataset, feature counts in each analyzed sample (*.xlsx)

***Additional file 2:*** Pilot Western Blotting experiment; results, materials and methods. **(\***.pdf)

***Additional file 3***: Proteomics dataset, quantified proteins and enrichment analysis (*.xlsx)

***Additional file 4:*** SynGO dataset, filtered gene list and enrichment analysis (*.xlsx)

## Declarations

### Ethics approval and consent to participate

The study protocols of the DeepCell project concerning the research on patient samples have been approved by the Ethics Committee of the Northern Ostrobothnia Hospital District (DeepCell, EETTMK:107/2016). All patient-derived samples for research were obtained based on voluntary participation in the study, and written informed consent was obtained from all patients or parents or guardians of such participating in this study under the guidance of a physician. When personal information on a patient is collected, stored, accessed and used, special attention was paid to the protection of the confidentiality of the subject according to the European Directive 95/46/EC.

### Consent for publication

Patients have given their consent for publication provided that they remain anonymous.

## Availability of data and materials

The processed RNA-seq and LC-MS datasets are described and available via the BioStudies database (https://www.ebi.ac.uk/biostudies/) under accession number S-BSST667. Raw data from the LC-MS analysis is accessible via Proteomics IDEntifications Database (PRIDE, https://www.ebi.ac.uk/pride/archive/projects/PXD026936) (30,31). The raw datasets generated by RNA sequencing during the current study are not publicly available, because of the risk that the data originated by sequencing-based technology may reveal enough variants to identify an individual. All the rest relevant data are supplied within the current publication.

## Competing interests

The authors declare no competing interests.

## Funding

This work was supported by the Academy of Finland [Decision numbers #311934 R.H. (profiling programme) and #331436 J.U.], Pediatric Research Foundation, Finland (J.U. and R.H.), Biocenter Oulu (J.U. and R.H.), Biocenter Finland, Special State Grants for Health Research, Oulu University Hospital, Finland (J.U.) and the Terttu Foundation, Oulu University Hospital, Finland (J.K.).

## Authors’ contributions

SK: study design, sample preparation protocols, sample collection, laboratory experiments, data analysis, and drafting the manuscript

JT: LC-MS experiments, method development and data analysis

ML: patient recruitment, sample collection, clinical data

AS: RNA-seq method development

BG: RNA-seq data analysis

PM: RNA-seq experiment design

JU (shared last): study design, research ethics, supervision and funding

MV (shared last): study design, supervision and funding, method development, resources for LC-MS analysis

JK (shared last): study design, sample collection and funding

RH (shared last): study design, sample preparation protocols, sample collection, laboratory experiments, drafting the manuscript, supervision and funding

All authors discussed the results, read, and approved the final manuscript.

## Acknowledgements

Laboratory technician Pirjo Keränen is acknowledged for her assistance in sample collection. LC-MS analysis was performed at the Proteomics unit, University of Helsinki, and RNA-seq experiments were performed by the Sequencing unit of Institute for Molecular Medicine Finland FIMM Technology Centre, University of Helsinki. Proteomics and Sequencing units are supported by Biocenter Finland.

## Notes

https://www.ebi.ac.uk/biostudies/studies/S-BSST667

https://www.ebi.ac.uk/pride/archive/projects/PXD026936

### Competing Interest Statement

The authors have declared no competing interest.

### Summary of Updates

The structure of the manuscript is changed. The text is updated and new references to publications and datasets are added.

