## Additional file 2 for "Analysis of human brain tissue derived from DBS surgery"

### SDS-PAGE and Western blotting

First, we analyzed one pilot sample (not included in mass spectrometry sample series) to evaluate the protein content and size distribution using SDS-page and western blot (Figure 1). Stain-free gel imaging revealed that the samples contain proteins of a large size range. Western blotting detected neuron- and glia-derived proteins in the samples. Western blotting was used to identify proteins in the samples that are expressed by nervous tissue –specific cell types such as neurofilament L (neuronal marker), GFAP (astrocyte marker) and CNPase (oligodendrocyte marker).

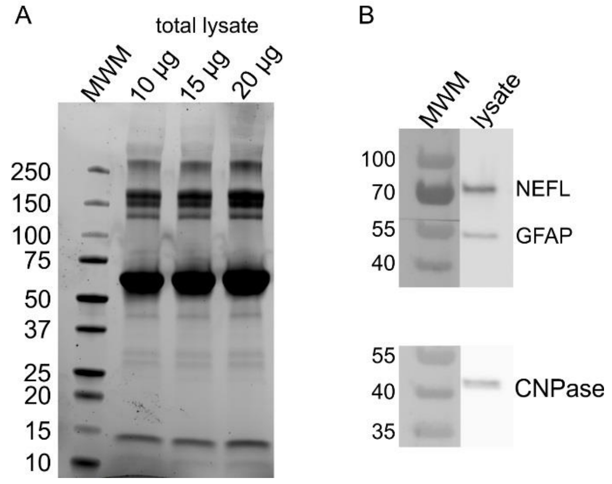


**Figure 1.** Image of SDS-PAGE gel and Western blot. A) 4-20% SDS-PAGE gradient gel (BioRad) was loaded with 10µg, 15 µg and 20µg of sample (total protein) and proteins were detected using stain-free ChemiDoc MP imaging system. B) Western blotting was used for identification of neurofilament L (neuronal marker, 1:1000), GFAP (astrocyte marker, 1:1000) and CNPase (oligodendrocyte marker, 1:1000).

### Methods

#### *Sample preparation for SDS-PAGE and Western blot*

The frozen tissue pellet was thawed on ice and the proteins were solubilized as follows: 25 µl of protein solubilization solution (phosphate buffered saline containing 1,5 % n-Dodecyl β-D-maltoside (Sigma-Aldrich) and Protease Inhibitor cocktail (Pierce)) was added onto the pellet and mixed by vortexing. The samples were incubated on ice for 40-50 minutes and vortexed occasionally. After the incubation, the samples were centrifuged for 20 minutes at 20 000 x g at +4°C. The supernatant containing the solubilized proteins was transferred into a new microcentrifuge tube. The protein concentration of the sample was determined by using Bradford assay (Pierce). Solubilized protein samples were prepared for SDS-PAGE by adding Laemmli sample buffer (Biorad) containing 2-mercaptoethanol and incubating at room temperature overnight prior to running SDS-PAGE. To evaluate the protein content and size distribution in the sample, 4-20% Mini-PROTEAN® Stain-free TGX™ Precast Protein Gel (BioRad) was loaded with 10µg, 15 µg and 20µg of sample (total protein) and proteins were detected using stain-free ChemiDoc MP imaging system. For Western blotting, 20 µg of the protein sample was loaded into the 4–20% Mini-PROTEAN® TGX™ Precast Protein Gel (Biorad).

The gel was blotted onto nitrocellulose membrane (Trans-Blot® Turbo™ Mini Nitrocellulose Transfer Pack, BioRad) with Trans-Blot Turbo transfer system (Biorad). The primary antibodies to detect GFAP, CNPase and Neurofilament-L were from Neuronal Marker IF Antibody Sampler Kit (#8572, Cell Signaling Technology, MA, United States). The secondary antibody used was Goat anti-Rabbit IgG horseradish peroxide (HRP) (Abcam, Cambridge, UK), dilution 1:10000. The bands were visualized using WesternBright ECL Spray substrate (Advansta, CA, United States).
